# Regulation of *PYR/PYL/RCAR* ABA receptors mRNA stability: involvement of miR5628 in decay of *PYL6* mRNA

**DOI:** 10.1101/2023.01.17.524441

**Authors:** João G. P. Vieira, Gustavo T. Duarte, Carlos H. Barrera-Rojas, Cleverson C. Matiolli, Américo J. C. Viana, Lucas E. D. Canesin, Renato Vicentini, Fabio T. S. Nogueira, Michel Vincentz

## Abstract

Hormone signaling fine-tuning involves feedback regulatory loops. Abscisic acid (ABA) plays key functions in development and tolerance to abiotic stress. ABA is sensed by the PYR/PYL/RCAR receptors and it also represses their gene expression. Conversely, ABA induces *PP2C* phosphatases expression, which are negative regulators of the ABA signaling pathway. This feedback regulatory scheme is likely important for the modulation of ABA signal transduction. Here, we provide a new insight into the mechanisms underlying the ABA-induced negative control of *PYR/PYL/RCAR* expression in *Arabidopsis thaliana*. The strong and sustained repression of *PYR/PYL/RCARs* revealed by ABA time course treatment defines the regulation of receptors genes as an important step in resetting the ABA signaling pathway. Transcription inhibition by cordycepin showed that destabilization of *PYL1/4/5/6* mRNA is involved in ABA-induced repression of these genes. Furthermore, genetic evidence indicated that decapping may play a role in *PYL4/5/6* mRNAs decay. In addition, we provide evidence that the *Arabidopsis-specific* microRNA5628 (miR5628), which is transiently induced by the ABA core signaling pathway, guides the cleavage of *PYL6* transcript in response to ABA. After cleavage, the resulting RISC 5’- and 3’-cleaved fragments of *PYL6* mRNA may be degraded by exoribonuclease XRN4. MiR5628 is an evolutionary novelty that may contribute, with decapping and XRN4 activities, to enhance *PYL6* mRNA degradation. Thus, control of stability of *PYR/PYL/RCAR* transcripts is an important step in maintaining homeostasis of ABA signaling.

**One Sentence Summary:** Attenuation of ABA signaling involves destabilization of *PYL1/4/5/6* transcripts. ABA core signaling induces miR5628 expression to enhance *PYL6* mRNA degradation in conjunction with decapping and XRN4 activities.

## INTRODUCTION

To adjust their physiology and growth in response to constant environmental changes, plants developed mechanisms to sense and transduce external and endogenous signals into adequate growth and developmental responses. This ability of a biological system to regulate its internal state in the face of external perturbations is known as homeostasis (Somvanshi et al., 2015). Control of homeostasis is the standard in hormone responses, whereby the signaling cascade must be dampened after an initial trigger to avoid overreactions. Such a control involves feedback loops. For instance, auxin, gibberellin, ethylene, strigolactone, cytokinin and brassinosteroids signaling are known to trigger the activation of negative regulators as part of the responses induced by these hormones (Teale et al., 2006; Davière and Achard, 2013; Zhu et al., 2013; Rai et al., 2015; Waters et al., 2017; Kieber and Schaller, 2018).

Similarly, the modulation of abscisic acid (ABA) signaling also involves the induction of negative regulators (*i.e., PP2C* phosphatases) and downregulation of positive regulators (*i.e*., *PYR/PYL/RCAR* ABA receptors), as part of the feedback loop for the attenuation of the hormone signal transduction (Song et al., 2016). Basal ABA levels, which range from 6 to 32 nM in *Arabidopsis thaliana*, support plant growth and development via a beneficial effect on water status of plants (Yoshida et al., 2019), while higher level of ABA plays an important role in responses to abiotic stress conditions such as drought, heat, cold and high salinity (Kavi Kishor et al., 2022). In *A. thaliana*, the ABA core signaling pathway is composed by 14 ABA receptors PYRABACTIN RESISTANCE 1 (PYR1)/PYR1-LIKE (PYL)/REGULATORY COMPONENT OF ABA RECEPTOR (PYR/PYL/RCAR), classified into three subgroups; nine clade A type 2C, PROTEIN PHOSPHATASES (PP2C); and three SNF1-related protein kinases 2 from subclass III (SnRK2) (Ma et al., 2009; Park et al., 2009). In the absence of ABA, PP2Cs continuously inhibit the signaling pathway by repressing SnRK2s activity (Ma et al., 2009; Park et al., 2009). The PYR/PYL/RCAR receptors of subgroup I have higher affinity for ABA than subgroups II and III, which respond to higher ABA levels (Tischer et al., 2017; Yoshida et al., 2019). Thus, subgroup I PYR/PYL/RCAR would be more involved in growth processes, while subgroups II and III would have a more prominent role under stress conditions, when ABA levels can increase up to 30-fold (Urano et al., 2017). In either case, ABA is perceived by PYR/PYL/RCARs, which inhibit the PP2Cs, thus releasing SnRK2 kinases from the negative regulation (Ma et al., 2009; Park et al., 2009). These kinases then can trigger changes in gene expression by activating ABA-responsive transcription factors and by modulating the activity of protein complexes involved in mRNA stability control (Wang et al., 2013).

The control of mRNA stability is an important mode of regulation of *PYR/PYL/RCAR* and *PP2C* expression (Wang et al., 2015; Wawer et al., 2018). In *A. thaliana*, cytoplasmic mRNA turnover pathway is initiated by poly(A) tail shortening. Next, the deadenylated mRNA may either be channeled into the 3’-5’ mRNA decay pathway, which involves degradation by the exosome, or be targeted by the 5’-3’ mRNA decay pathway, which consists of the DECAPPING 2 (DCP2) enzyme and its activators DCP1, DCP5, VARICOSE, and PAT1. Finally, 5’-decapped transcripts are degraded by the exoribonuclease XRN4 (Chantarachot and Bailey-Serres, 2018). This mRNA turnover pathway is conserved among eukaryotes (Mugridge et al., 2018). Alternatively, transcripts degradation can be triggered by microRNAs (miRNAs) pathways. MiRNAs biogenesis begins with the transcription of MIR genes, by RNA polymerase II, generating a primary transcript (pri-miRNA). This transcript generates a hairpin-shaped secondary structure that is processed by DICER-LIKE1 (DCL1) generating a miRNA-miRNA duplex. This duplex is methylated by HEN1 to be transported from the nucleus to the cytoplasm. One of the strands of the duplex is incorporated into the Interfering RNA Silencing Complex (RISC). In the cytoplasm, the miRNA will recognize its target by base complementarity, cleaving it by Argonaut, or repressing translation (Barrera-Rojas et al., 2021).

Our previous observation that ABA and glucose act in synergy to destabilize the mRNA of the transcription factor *bZIP63* (Matiolli et al., 2011), prompted us to investigate the impact of ABA on the control of mRNA stability at genomic scale. Interestingly, we noticed that ABA promotes the destabilization of *PYL4, PYL5* and *PYL6* receptors transcripts. Here, we confirm that ABA accelerates the decay of these transcripts. We provide evidence that the ABA core signaling pathway induces the *Arabidopsis-specific* miR5628 expression in response to ABA, which in turn promotes *PYL6* mRNA cleavage and mRNA decay in a XRN4-dependent way. Moreover, decapping was also found to be involved in the control of *PYL4/5/6* receptors mRNA decay in response to ABA. The control of the stability of *PYR/PYL/RCAR* transcripts is proposed to be part of a feedback regulatory loop participating in the attenuation and resetting of the ABA signaling.

## RESULTS

### Negative feedback of ABA signaling involves sustained repression of *PYR/PYL/RCAR* gene expression

Regulatory feedback acting upon the ABA signaling pathway has been described (Song et al., 2016). To obtain new insight into the underpinning mechanism, we set-up an experimental design to evaluate the response of the ABA core signaling pathway genes in response to 1 μM ABA, which is likely to mimic abiotic stress conditions (Tischer et al., 2017; Urano et al., 2017). We have analyzed the expression profile of seven of the fourteen representative members of *PYR/PYL/RCAR* receptors multigene family (Supplemental Fig. S1A). We also have measured the expression of six representative members of *PP2C* phosphatases and all three *SnRK2* kinases from subclass III (Supplemental Fig. S1A). After one-hour of ABA treatment, the expression of all seven *PYR/PYL/RCARs* were found to be repressed in response to the hormone (Supplemental Fig. S1B). In contrast, the six *PP2Cs* phosphatases and *SnRK2.6* were induced by ABA, while the other two members of the subclass III *SnRK2* kinases did not respond (Supplemental Fig. S1B). Thus, our experimental conditions recapitulate the ABA-induced regulation of ABA core signaling genes described earlier (Song et al., 2016).

To further explore the dynamic regulation of this representative subset of the ABA core signaling genes (Supplemental Fig. S1), we have analyzed the time course changes of their mRNA profiles in response to prolonged (16 hours) and transient (30 minutes) ABA treatments. These treatments were meant to mimic ABA-mediated responses to variable durations of stress exposure. ABA-treatment for 16 h resulted in a continuous repression of all ABA receptors analyzed but *PYL1* (Fig. 1A; Supplemental Fig. S2), while all *PP2Cs* and *SnRK2.6* genes were transiently induced between one and four hours after the beginning of ABA treatment, followed by a gradual decrease and stabilization of their mRNA levels (Fig. 1A; Supplemental Fig S2). *SnRK2.2* and *SnRK2.3* were induced later in comparison to the other ABA core signaling elements (Fig. 1A). The continuous repression of most *PYR/PYL/RCAR* in comparison to the transient induction of the *PP2C* genes suggests that the negative regulatory feedback acting on the control of *PYR/PYL/RCAR* expression has an important role in attenuating the ABA signaling pathway. The induction of *SnRK2.2/3/6* by ABA possibly reflects a way to amplify ABA-mediated stress responses.

**Figure 1:**
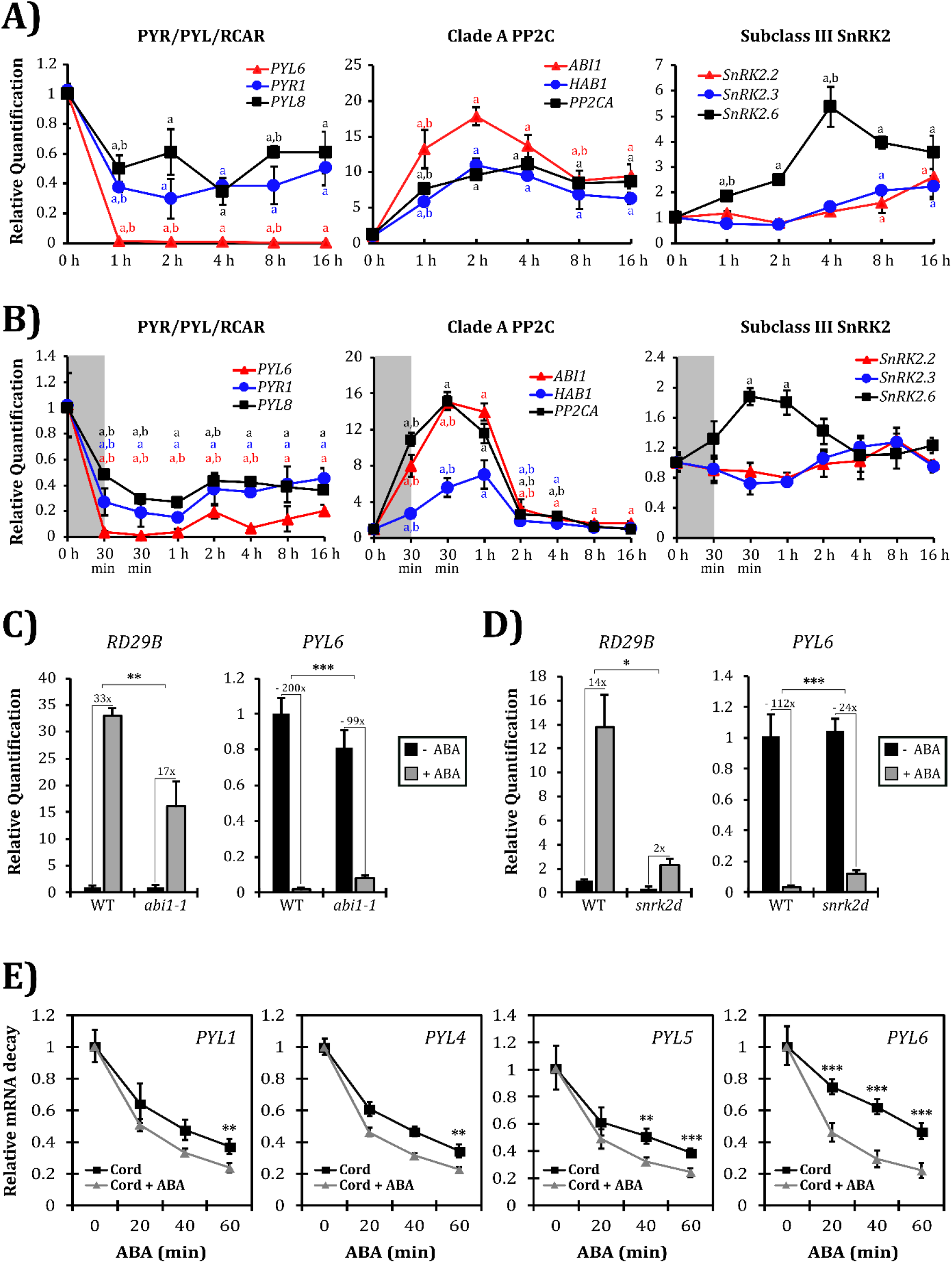
Regulation of the expression of ABA core signaling pathway genes by ABA. (**A**) mRNA profile of members of the ABA core signaling pathway in response to long-term ABA treatment. One member of each subgroup of PYR/PYL/RCAR and three members of clade A PP2Cs (Supplemental Fig. S1A) are represented, along with subclass III SnRK2 members. An expanded analysis with *PYR/PYL/RCAR* and *PP2C* genes is given in Supplemental Fig. S2. Seedlings were treated with 1 μM ABA for up to 16 hours. (**B**) mRNA profile of members of the ABA core signaling pathway in response to transient ABA-treatment. Seedlings were treated with 1 μM ABA for 30 min, then ABA was removed from the medium. Sampling was performed before the treatment (0 h), at the end of the 30 min (in the gray box) of ABA-treatment, and at different time points after hormone removal. An expanded analysis with *PYR/PYL/RCAR* and *PP2C* genes is given in Supplemental Fig. S3. In panels A and B “a” means significant difference between each time point vs the untreated control and “b” each time point vs. the previous one. Responses were significantly different for fold changes in mRNA levels ≥ |1.5| and p < 0.05, according to Student’s t-test. The color of the letter refers to the gene evaluated. (**C**) The negative feedback regulation of the ABA core signaling pathway requires ABA-signalization. Wild type (WT) and *abi1-1* seedlings were treated with 1 μM ABA for 1 hour before sampling. Expression of the ABA-induced *Rd29B* gene was used as positive control of ABA-promoted responses. *PYL6* response is shown and a complete analysis of the expression of six other representative ABA receptors is provided in Supplemental Fig. S4. The expression levels are given in comparison to the untreated WT. (**D**)Kinases SnRK2 are involved in the negative feedback of *PYR/PYL/RCAR* genes in response to ABA. WT and the double kinase mutant *snrk2.2/snr2.3 (snrk2d*) seedlings were treated with 1 μM ABA for 1 hour before sampling. Expression of the ABA-induced *Rd29B* gene was used as positive control of ABA-promoted responses. *PYL6* response is shown and a complete analysis of the expression of the six other representative ABA receptors is provided in Supplemental Fig. S6. The expression levels are given in comparison to the untreated WT. (**E**) ABA regulates *PYL1, PYL4, PYL5* and *PYL6* mRNA stability. To evaluate ABA-mediated control of mRNA stability, seedlings were pre-treated for 1 h with 100 μM cordycepin (Cord) to inhibit transcription, followed by the addition of ABA 1 μM (Cord + ABA). After cordycepin pre-treatment sampling was performed at 20, 40 and 60 minutes with and without ABA. Relative expression values in Cord + ABA condition that are lower than the respective Cord treatment alone indicate that the stability of the transcript was decreased in response to ABA. A complete analysis of the mRNA stability of representative *PYR/PYL/RCAR* genes is given in Supplemental Table S3. For all experiments, values are the mean of five biological replicates ± standard deviation. Responses were significantly different for fold changes in mRNA levels ≥ |1.5| and p < 0.05, according to Student’s t-test (* < 0.05; ** < 0.005; *** < 0.0005).

The changes of ABA core signaling genes expression in response to short term ABA application should give clues about how the pathway is reset to its steady state levels (*i.e*., how it recovers the original transcript amounts prior to ABA treatment). Therefore, we have performed a transient ABA treatment for 30 min and monitored the mRNA levels of the representative subset of ABA core signaling pathway genes for 16 h following removal of the hormone (Fig. 1B; Supplemental Fig. S3). The maximum response of all evaluated ABA core signaling transcripts occurred between the end of the 30 min of ABA treatment and 1 h after the hormone removal (Fig. 1B; Supplemental Fig. S3). The only exceptions were *SnRK2.2* and *SnRK2.3*, which were not affected (Fig. 1B). Within a time frame of eight hours after the transitory ABA treatment, *HAB1, PP2CA, HAI2* PP2Cs and *SnRK2.6* transcripts recovered their original levels, and *ABI1, ABI2* and *HAI1* transcripts clearly tend also to do so (Fig. 1B; Supplemental Fig. S3). On the other hand, the representative receptors had a slightly different behavior, *PYR1, PYL1/4/5/6* and *PYL8* showed a slow recovery rate of their original transcript levels, reaching at most 60% their initial levels (Fig. 1B; Supplemental Fig. S3), while *PYL2* has not shown a tendency to recover over this time period (Supplemental Fig. S3). These results suggest that the control of ABA receptors gene expression is an important aspect of the reset process of the ABA signaling pathway.

To address whether a functional ABA core signaling pathway is required to trigger the feedback downregulation of the gene expression of ABA receptors, we have adopted two approaches. First, we compared the expression of *PYR/PYL/RCAR* genes in response to ABA between the wild-type (WT) and the dominant *abi1-1* mutant, which maintains SnRK2s dephosphorylated and, therefore, is insensitive to ABA (Umezawa et al., 2009). *Rd29B* is a readout gene for ABA signaling (Yoshida et al., 2015) and was found to be two times less effectively induced by ABA in *abi1-1* than in the WT (Fig. 1C). ABA-promoted repression of *PYL6* (Fig. 1C) and *PYR1, PYL2, PYL4, PYL5, PYL6* and *PYL8* receptor genes is attenuated in the *abi1-1* mutant in comparison to the WT (Supplemental Fig. S4).

The second approach consisted in evaluating the participation of subclass III SnRK2 kinases in the negative feedback regulation of ABA receptors by global kinase inhibition with staurosporine and by monitoring the SnRK2 double-mutant, *snrk2.2/snrk2.3* (*snrk2d*), in which ABA responses are attenuated (Fujii et al., 2007). Staurosporine was effective in reducing the induction of *Rd29B* by ABA, indicating that inhibition of subclass III SnRK2 kinases was partially achieved (Supplemental Fig. S5). In the presence of staurosporine, the repression of *PYL1, PYL2, PYL4, PYL5, PYL6* and *PYL8* by ABA was significantly attenuated as compared to the control (Supplemental Fig. S5). In addition, a weaker induction of the *ABI1, ABI2, HAI1* and *HAI2* phosphatase genes by ABA in the presence of staurosporine was detected (Supplemental Fig. S5). The results suggest that in response to ABA, subclass III SnRK2 kinases are required for efficient repression and induction of *PYR/PYL/RCAR* and of clade A *PP2C* genes, respectively. As expected, induction of *RD29B* by ABA was significantly reduced in *snrk2d* (Fig. 1D). The ABA-promoted repression of *PYR1, PYL2, PYL4, PYL6* and *PYL8* receptors is attenuated in *snrk2d* mutant in comparison to the WT (Fig. 1D; Supplemental Fig. S6), while *PYL1* and *PYL5* showed a similar tendency (Supplemental Fig. S6). ABA-based *PYL6* repression was more strongly attenuated in the *snrk2d* compared to the others receptor genes (Fig. 1D). These results further support the participation of SnRK2 in triggering repression of most *PYR/PYL/RCAR* in response to ABA. Together, these data suggest the ABA-promoted *PYR/PYL/RCAR* repression relies on a functional ABA signalosome.

Since *PYR/PYL/RCARs* emerge as key targets of the negative regulatory feedback and resetting of the ABA signaling pathway, the mechanisms underlying the repression of ABA receptors genes were further investigated. To distinguish transcriptional from mRNA decay regulations, we blocked transcription with cordycepin and analyzed changes of receptors mRNA levels. Since ABA-induced expression of *Rd29B* is known to be mainly transcriptional, this gene was, therefore, used as a control to monitor the efficiency of transcription inhibition by cordycepin (Matiolli et al., 2011). ABA-induced up-regulation of *Rd29B* mRNA was found to be reduced by 93% by cordycepin, indicating that transcriptional inhibition was efficient (Supplemental Table S3). *PYL8*(subgroup I), *PYL4, PYL5, PYL6* (subgroup II), *PYR1, PYL1* and *PYL2* (subgroup III) transcripts half-life were reduced by ABA to different extents (Supplemental Table S3). ABA-induced decay of *PYL4, PYL5* and *PYL6* mRNAs (subgroup II) was more accentuated than decay of receptors transcripts from subgroup I and III (Fig. 1E; Supplemental Table S3 and Supplemental Fig. S7). *PYL6* mRNA is the most strongly destabilized transcript with a half-life reduction around 50% (Fig. 1E, Supplemental Table S3). Together, these observations indicate that the ABA-induced downregulation of *PYR/PYL/RCAR* members expression relies, at least partly, on the control of the stability of their mRNAs.

Interestingly, *PYL6* was the only receptor for which ABA-promoted transcript levels reduction was more pronounced in the absence of cordycepin (200-fold reduction by ABA *versus* 77-fold reduction by ABA + cordycepin; Supplemental Table S3). This result raised the possibility that ABA-induced *PYL6* mRNA decay requires transcriptional induction by ABA of a regulatory factor that modulates *PYL6* transcript stability. The hypersensitivity to ABA of different mutants affecting the miRNAs pathway (Duarte et al., 2013) prompted us to examine the possibility that a miRNA, whose expression would be induced by ABA, could be involved in *PYL6* mRNA destabilization.

### MiR5628 is involved in the control of *PYL6* mRNA stability in response to ABA

To evaluate the involvement of the miRNA pathway in the control of *PYL6* mRNA stability, we analyzed its expression in mutants defective in miRNA biogenesis (*hyl1-2* and *se-1*) and activity (*ago1-25*). After ABA treatment, *PYL6* mRNA levels were 2.2, 1.7 and 2.2-fold higher in *hyl1-2, se-1* and *ago1-25*, respectively, when compared to the WT (Supplemental Fig. S8). These results support the hypothesis that the miRNA pathway is involved in the control of *PYL6* transcript stability.

Based on *in silico* prediction analyzes (psRNA-Target software), four putative miRNAs which could recognize *PYL6* transcript were identified. MiR8175 could bind to *PYL6* coding sequence, whereas miR5628, miR5021 and miR840-3p, could target the 3’UTR region of the transcript (Fig. 2A; Supplemental Fig. S9). 5’-RACE analyzes were carried out to investigate whether these four miRNAs can guide the cleavage of *PYL6* transcript. *SPL9*, which is a target of the conserved miR156 (Wang et al., 2009), was used as positive control for ligation of the GeneRacer RNA oligo to the RISC 3’-cleaved fragment (Supplemental Fig. S10A). After 30 minutes of ABA treatment, 5’-RACE products were obtained for miR5628 (Supplemental Fig. S10A), but not for the other three miRNAs. The 5’-RACE PCR product related to potential miR5628-guided cleavage were cloned and sequenced. Three cloned 5’-RACE products were found to match position 11 and one clone matched position 15 of miR5628 site in *PYL6* mRNA, suggesting that, indeed, miR5628 guides *PYL6* mRNA cleavage (Fig. 2A). The 5’-end of the remaining 43 sequenced clones matched sequences downstream to miR5628 recognition site (Fig. 2A) and may represent 5’-end products of XRN4 exoribonuclease degradation. This hypothesis was supported by the observation that, after ABA treatment, *PYL6* mRNA 3’UTR region accumulated more in *xrn4-5* mutant than in the WT (Supplemental Fig. S10B). These results suggest that miR5628 promotes cleavage of *PYL6* mRNA in the 3’UTR region and XRN4 degrades the RISC 3’-cleaved fragment of *PYL6* mRNA.

**Figure 2:**
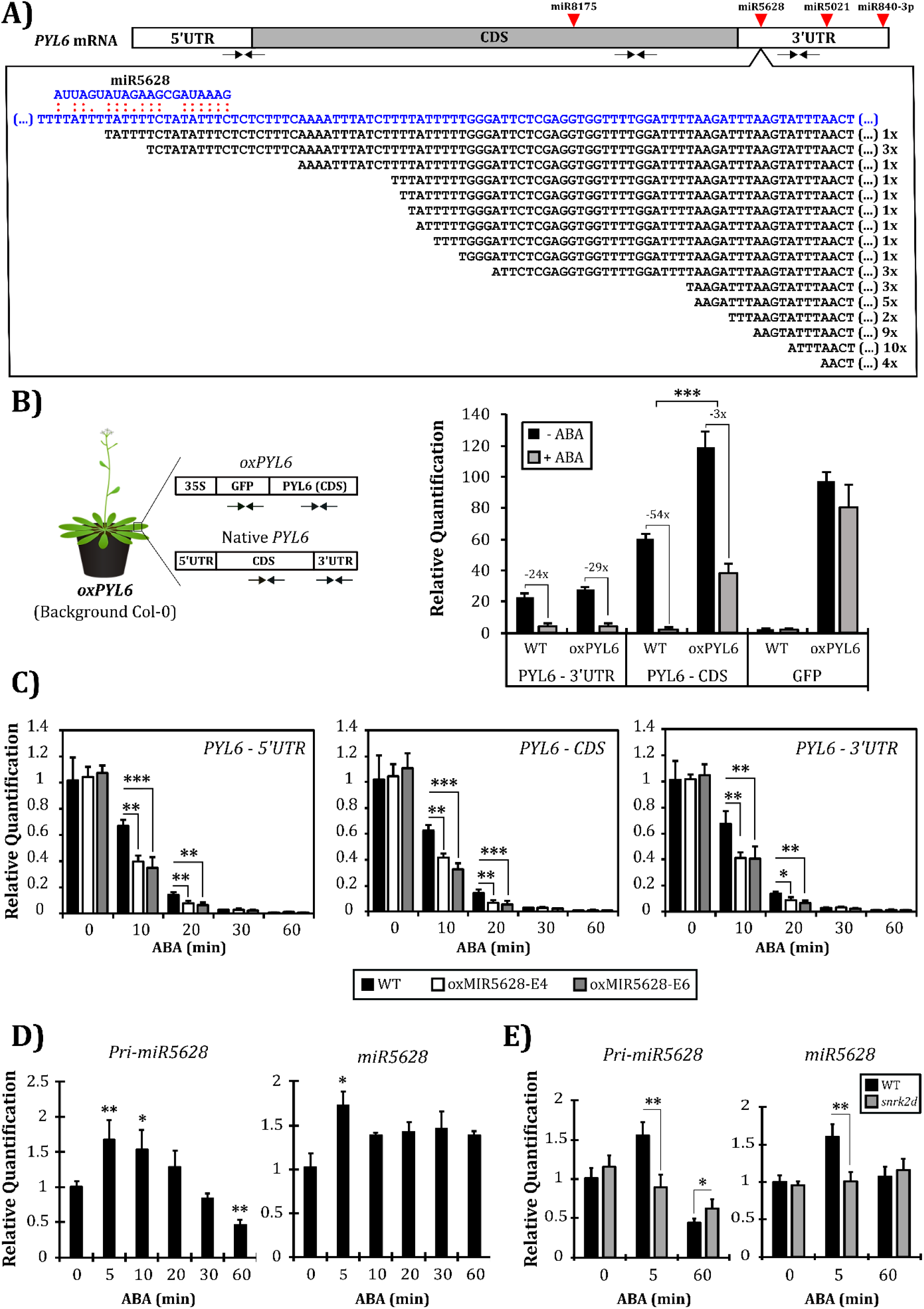
MiR5628 promotes cleavage of *PYL6* mRNA in response to ABA. (**A**) Schematic representation of *PYL6* mRNA with the putative miR8175, miR5628, miR5021 and miR840-3p and the position of their binding sites in *PYL6* mRNA. 5’-RACE analysis of *PYL6* mRNA cleavage by miR5628 in Col-0. Forty-seven cloned sequences (in black) mapped to the miR5628 recognition site (position 784-804 bp of *PYL6* mRNA) or downstream of it in the 3’UTR sequence (blue). The number of occurrences of each sequence is indicated. (**B**) The 3’UTR of *PYL6* transcript is required for proper control of its stability in response to ABA. The relative amounts of the CDS and 3’UTR sequences of *PYL6* mRNA was quantified in wild type (WT) and in a transgenic line expressing the 35S:GFP:PYL6 (Col-0 background), which lacks the *PYL6* 3’UTR region (*oxPYL6*). Amplification of *GFP* sequence allows to quantify specifically the *oxPYL6* fusion. Fourteen days old seedlings grown under continuous light were treated with 1 μM ABA for 30 min. This analyze is representative of three independent experiments. (**C**) miR5628 participates in the degradation of *PYL6* transcript in response to ABA. The amount of *PYL6*-5’UTR, CDS and 3’UTR sequences (primers positions are shown in panel A) was quantified at different time points of a 60 min treatment with 1 μM ABA in samples of oxMIR5628 (E4 and E6) and WT seedlings. Relative expression values of each genotype were obtained in comparison to the untreated WT. This analyze is representative of two independent experiments. (**D**) ABA transiently induces both the *pri-miR5628* and the mature *miR5628*. Col-0 seedlings were treated with 1 μM ABA and sampling was performed before (0 min) and at 5, 10, 20, 30 and 60 minutes after ABA application. Relative expression values were obtained in comparison to the untreated condition (time point of 0 min). This analyze is representative of four independent experiments. (**E**) A functional ABA signaling is required to transient induction of *pri-miR5628* and the mature *miR5628* in response to ABA. WT and the double kinase mutant *snrk2d (snrk2.2/snr2.3*) seedlings were treated with 1 μM ABA and sampling was performed before (0 min), at 5 and 60 minutes after ABA application. The relative expression levels are given in comparison to the untreated WT. In panels B, C, D and E, values are the means of five biological replicates ± standard deviation. RT-qPCR was carried out to gene quantification. Responses were significantly different for fold changes in mRNA levels ≥ |1.5| and p < 0.05, according to Student’s t-test (* < 0.05; ** < 0.005; *** < 0.0005).

Additionally, we analyzed the expression of a transgenic line (Col-0 background) expressing a fusion between *Green Fluorescent Protein* (GFP) and the coding sequence of *PYL6* under the control of the 35S promoter but lacking the *PYL6*-3’UTR sequence (*oxPYL6*), which contains the miR5628 target site (Fig. 2B). Quantification of endogenous *PYL6* mRNA using primers covering its 3’UTR showed that in both WT and *oxPYL6*, the native *PYL6* transcripts are equally downregulated by ABA (Fig. 2B). Using *PYL6* coding sequence (CDS)-specific primers, two-times more *PYL6*-CDS mRNA sequences were detected in *oxPYL6* than in the WT because the *PYL6*-CDS-specific primers amplify the endogenous *PYL6* and the *GFP:PYL6* fusion transcripts (Fig. 2B). Yet, ABA treatment promoted only a three-fold reduction of *PYL6*-CDS mRNA in *oxPYL6* as compared to a 54-fold repression in the WT (Figure 2B). Since promoter 35S is not regulated by ABA (Chu and Jeng, 2002), this difference is not due to ABA-mediated transcriptional effect and could, therefore, be a consequence of the inability of mRNA decay regulations to act on the *GFP:PYL6* fusion transcript which lacks the 3’UTR region (Fig. 2B). This possibility is supported by the observation that the GFP-mRNA sequence is only marginally repressed by ABA (Fig. 2B). Taken together, these results support the notion that *PYL6* repression by ABA rely, at least in part, on the 3’UTR sequence, which contain the miR5628-target sequence.

To get further confirmation of miR5628 involvement in *PYL6*-3’UTR cleavage, we used a dual-luciferase-based miRNA sensor system to evaluate whether miR5628 can guide *PYL6* mRNA cleavage *in vivo*. We made three different constructs based on pGreen_dualluc_3’-UTR sensor vector containing either the WT *PYL6* mRNA recognition site of miR5628, or a fully complementarity sequence to the miR5628 sequence (positive control). A negative control consisting of a non-complementary sequence of miR5628 was included (Supplemental Fig. S10C). *Nicotiana benthamiana* leaves were co-transformed with the reporter constructs. The native and positive control constructions showed a tendency to be downregulated in comparison to negative control further seggesting that miR5628 guides the cleavage of *PYL6* mRNA (Supplemental Fig. S10D). We also generated two transgenic lines overexpressing miR5628 (oxMIR5628-E4 and oxMIR5628-E6) (Supplemental Fig. S11) and analyzed the profile of accumulation of the *PYL6* (5’UTR, CDS and 3’UTR regions) along a time course of ABA treatment (Fig. 2C). In the absence of ABA there is no difference in the abundance of *PYL6* mRNA in the oxMIR5628-E4 and oxMIR5628-E6 lines compared to WT (Fig. 2C). However, all regions of *PYL6* transcript are significantly less abundant in the two oxMIR5628 lines compared to WT in the first 20 min of ABA treatment (Fig. 2C), while after 30 and 60 minutes, no difference in *PYL6* mRNA level was detected between oxMIR5628 and the WT (Fig. 2C). This result supports the hypothesis that miR5628 cleaves *PYL6* transcript and promotes its instability in an ABA-dependent manner.

Finally, 5’-RACE analysis was performed with mRNA sampled at 20 min of ABA treatment of oxMIR5628 lines to evaluate the cleavage-activity of miR5628 in these lines (Supplemental Fig S12). 166 cloned 5’-RACE products (52 for WT, 54 for oxMIR5628-E4 and 60 for oxMIR5628-E6) were found to match either the miR5628 recognition site in *PYL6* mRNA or downstream to it, but none was localized upstream to miR5628 recognition site (Supplemental Fig S12). This result further corroborates that miR5628 is involved in *PYL6* transcript decay by promoting *PYL6* transcript cleavage.

Based on transcriptional repression by cordycepin we raised the hypothesis that an efficient ABA-induced decay of *PYL6* mRNA rely on an ABA-inducible regulatory factor (Fig. 1E; Supplemental Table S3). MiR5628 emerged as a candidate (Fig. 2A-C), thus, its expression is expected to be regulated by ABA. Indeed, ABA treatment resulted in a transient induction of both *pri-miR5628* and mature *miR5628* (Fig. 2D), reinforcing the regulatory input of miR5628 on *PYL6* transcript stability. Moreover, this transient induction is completely abolished in the double kinase mutant *snrk2d* in response to ABA (Fig. 2E), suggesting that a functional ABA core signaling is required. Additionally, we analyzed the production of *pri-miR5628* and the mature *miR5628* in the miRNAs biogenesis mutants *hyl1-2* and *se-1*. As expected, the *pri-miR5628* accumulated more in both mutants compared to WT in response to ABA, while the mature miR5628 is less abundant in the mutant *hyl1-2* treated with ABA compared to WT and in *se-1* miR5628 is less abundant both in the presence and absence of ABA in relation to WT (Supplemental Fig. S13). These results indicate that ABA controls miR5628 biogenesis.

### oxMIR5628 affects ABA-induced responses

*PYL6* is more expressed during seed germination suggesting it plays a role in this developmental phase (Klepikova et al., 2016). Thus, we hypothesized that oxMIR5628 lines would be hyposensitive to ABA during germination (*i.e*., radicle emergence). The germination rate of oxMIR5628 genotypes was found to be higher at the concentration of 0.5 and 0.75 μM of ABA compared to WT, with the largest difference observed at 0.75 μM of ABA (Fig. 3A). At this concentration, the germination rate of oxMIR5628 E4 and E6 was 82% and 84%, respectively, while WT was 58% (Fig. 3A). These data confirm that oxMIR5628 lines are hyposensitive to ABA suggesting that miR5628 may impact ABA signaling during germination.

**Figure 3:**
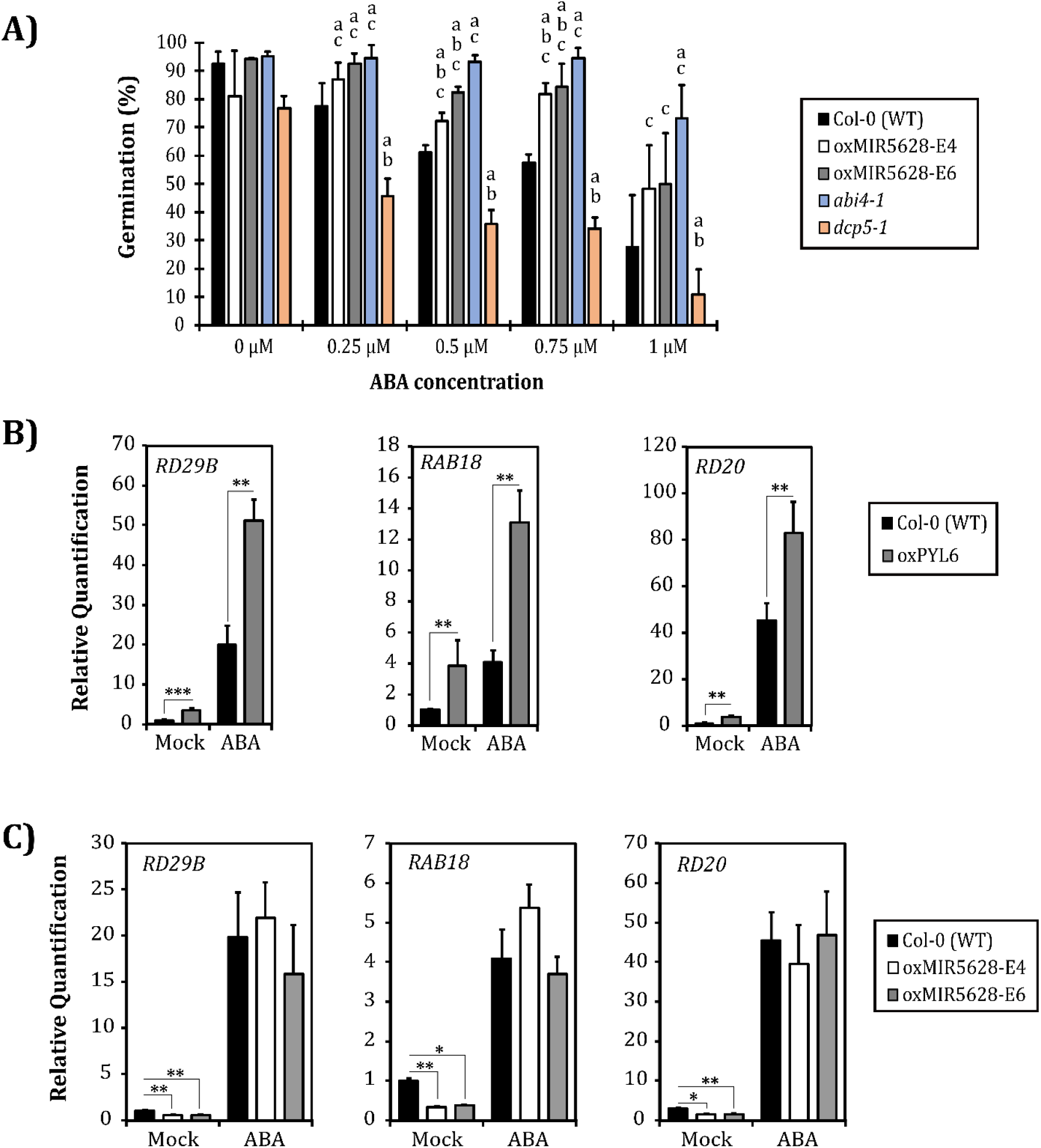
miR5628 impacts ABA signaling. (**A**) oxMIR5628 lines are hyposensitive to ABA during germination. The germination rate of WT, oxMIR5628 lines, *abi4-1* and *dcp5-1* genotypes were evaluated in continuous light on solid (0.25% of agar) MS/2 medium along a range of ABA concentrations (0, 0.25, 0.5, 0.75 e 1μM of ABA). The data represent the average of three biological replicates (three plates) with at least 30 seeds for each genotype. Seeds germination was monitored during six days after exposure to light. The seed germination rate (%) corresponds to the number of seeds that germinated each day for the total of seeds germinated at the sixth day. The graphic represents the differences observed at the second day. The mutants *abi4-1* and *dcp5-1* were used as control for ABA-hypo and - hypersensitive phenotypes, respectively. Responses were significantly different for changes in the germination rate according to Tukey and Scott-Knott test, which is represented by the letters “a” (statistical difference in relation to WT), “b” (statistical difference in relation to *abi4-1*) and “c” (statistical difference in relation to *dcp5-1*). This analyze is representative of two independent experiments. (**B**) ABA signaling readout genes (*RD29B, RAB18* and *RD20*) are upregulated in the *oxPYL6* genotype both in the presence and absence of ABA. WT and *oxPYL6* seedlings were treated with 1 μM ABA for 1 hour before sampling. Relative expression values of each genotype were obtained in comparison to the untreated WT. This analyze is representative of three independent experiments. **C**) ABA signaling readout genes (*RD29B, RAB18* and *RD20*) are downregulated in the oxMIR5628 lines. WT and oxMIR5628 (E4 and E6) seedlings were treated with 1 μM ABA for 1 hour before sampling. Relative expression values of each genotype were obtained in comparison to the untreated WT. This analyze is representative of three independent experiments. In panels B and C, values are the means of five biological replicates ± standard deviation. RT-qPCR was carried out to gene quantification. Responses were considered to be significantly different for fold changes in mRNA levels ≥ |1.5| and p < 0.05, according to Student’s t-test (* < 0.05; ** < 0.005; *** < 0.0005).

The impact of *PYL6* on ABA signaling can also be evaluated through expression analysis of ABA-responsive genes such as *RD29B, RD20* and *RAB18* in *oxPYL6* (Fujita et al., 2009; Yoshida et al., 2015). *RD29B, RD20* and *RAB18* genes were induced in the *oxPYL6* genotype compared to WT in the absence and presence of ABA (Fig. 3B), suggesting that *PYL6* participates in the control of expression of these ABA readouts genes. We then asked whether miR5628 overexpression could alter ABA-mediated induction of *RD29B, RD20* and *RAB18* as would be expected from miR5628-promoted *PYL6* mRNA degradation. In the presence of ABA, no significative difference in the expression of *RD29B, RD20*, or *RAB18* between oxMIR5628 genotypes and WT was detected (Fig. 3C). However, in the absence of ABA, these genes were downregulated in both oxMIR5628 genotypes compared to WT (Fig. 3C). *In silico* prediction analyzes (psRNA-Target software) showed that *RD29B, RD20* and *RAB18* mRNA are not direct target of miR5628. These results suggest that miR5628 indirectly controls the basal expression of ABA signaling readouts possibly by downregulating *PYL6* expression.

### MiR5628 has emerged in *A. thaliana*

We have analyzed whether miR5628 could recognize other *A. thaliana PYR/PYL/RCAR* mRNAs. Using the psRNA-Target tool only *PYL6* mRNA was found to be target of miR5628, suggesting a highly specific regulation (Supplemental Fig. S14). Indeed, none of the *PYR/PYL/RCAR* transcript evaluated but *PYL6* was downregulated in the oxMIR5628 lines compared to WT (Fig. 2C; Supplemental Fig. S15). Then, we have evaluated the evolutionary conservation of miR5628. To this end, BLAST searches were performed for precursor and mature miR5628 sequence similarity against plant genome databases (NCBI). Although a miR5628-related sequence was detected in *Brassica rapa*, the predicted secondary structure of this putative miR5628 precursor was unable to form a hairpin-loop, neither miR5628* and nor expression evidence of this locus was obtained. Finally, we analyzed the global miR5628 expression available in the *Arabidopsis Small RNA Database* (http://ipf.sustech.edu.cn/pub/asrd/) (Feng et al., 2020). MiR5628 is poorly expressed (maximum of 3 transcripts per million) as compared to conserved miRNAs (*e.g*., miR156a-3p with 3,808 transcripts per million) (Supplemental Table S4). The expression of miR5628 is quite comparable to other newly evolved miRNA such as miR5657, miR779.1 and miR865-3p (Supplemental Table S4), which is expected since lineage-specific miRNAs tend to be barely expressed (Fahlgren et al., 2012; Axtell, 2013; Hajieghrari and Farrokhi, 2021). Taken together, these results suggest that miR5628 may have evolved recently in *A. thaliana* lineage.

### Dynamic of *PYL6* mRNA decay

MiRNAs-mediated mRNA cleavage implies that exosome and XNR4 mediate the degradation of the RISC 5’- and 3’-cleaved fragments, respectively (Chantarachot and Bailey-Serres, 2018). Since miR5628 guides the cleavage of *PYL6* mRNA 3’UTR, a faster decay of this region in comparison to the 5’UTR and coding sequence (CDS) would be expected after ABA treatment. To address this possibility, a short-term kinetic (5 to 60 min) of ABA treatment was performed to monitor the rate of decay of different regions of *PYL6* transcript ranging from the 5’ to the 3’-end using region-specific primers (Fig. 4A). The kinetics degradation profiles of the three different parts of *PYL6* mRNA were found to be similar, suggesting that all parts of *PYL6* transcript are degraded synchronously (Fig. 4A). This conclusion raises the possibility that in addition to miR5628 activity, other mechanisms of mRNA decay are involved in *PYL6* transcript degradation. For instance, decapping has been associated to the control of *PYR/PYL/RCAR* transcript stability and mutants of this machinery are hypersensitive to ABA (Wawer et al., 2018). Additionally, RISC 5’-cleaved fragments of miRNA-targets were suggested to be degraded by the exoribonuclease XRN4 after decapping (Ren et al., 2014). Therefore, we have tested whether decapping followed by XRN4 activity could be involved in the degradation of *PYL6* mRNA.

**Figure 4:**
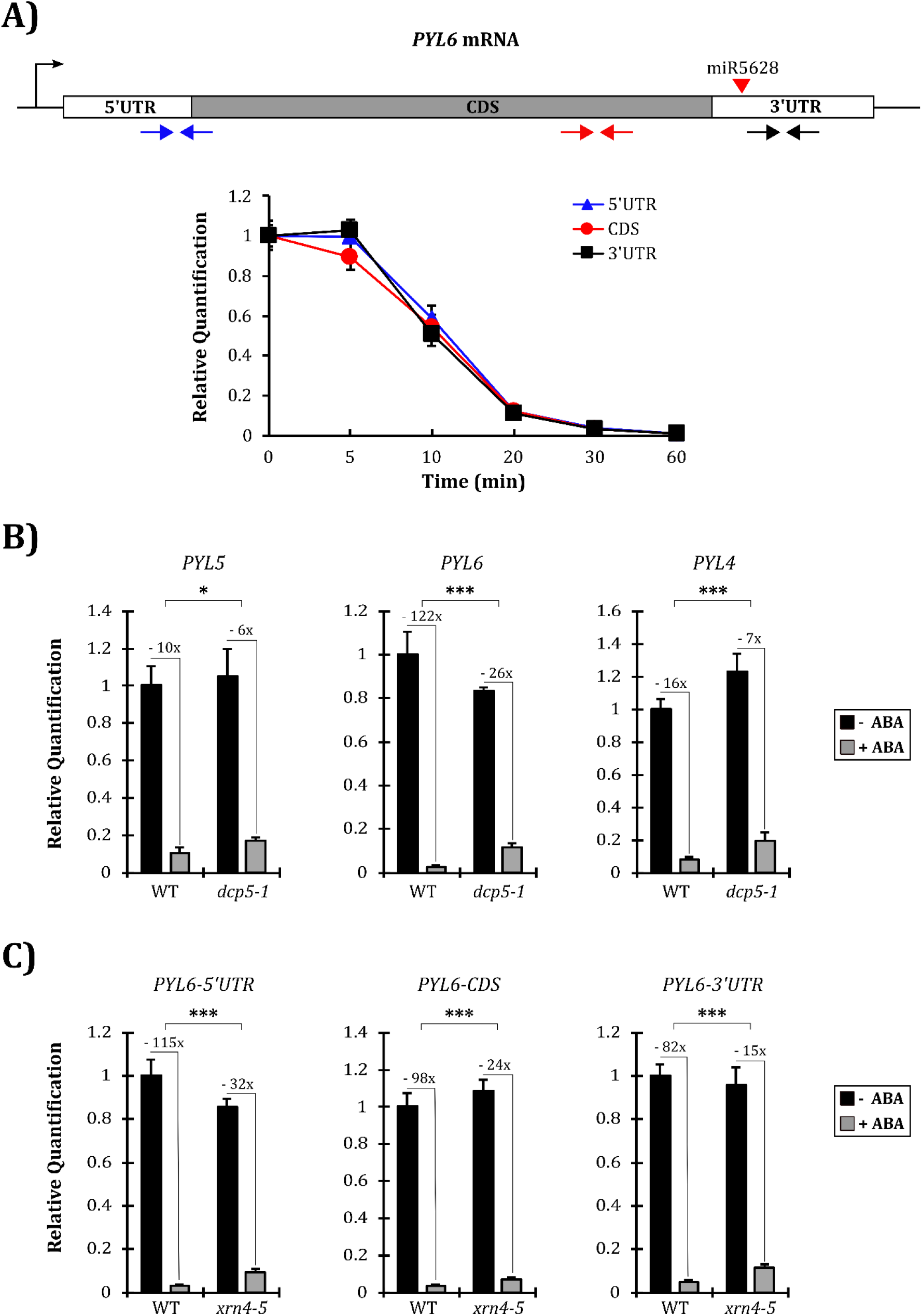
5’-3’ mRNA decay pathway is involved in the control of *PYL6* mRNA accumulation in response to ABA. (**A**) Schematic representation of *PYL6* mRNA with the 5’- and 3’-UTRs and coding sequence (CDS) regions, and miR5628 target site. The position of primer pairs that were used to measure the level of mRNA corresponding to different parts of *PYL6* transcript are shown. The different regions of *PYL6* were quantified by RT-qPCR along a time course of ABA treatment. Seedlings were grown for six days and were treated with 1 μM ABA. Sampling was performed before (0 min) and at 5, 10, 20, 30 and 60 min after ABA application. This analyze is representative of three independent experiments. (**B**) *PYL4, PYL5* and *PYL6* transcripts profiles were compared between the mutant *dcp5-1*, which is defective in decapping activity, and the wild type (WT) in response to ABA treatment. (**C**) Accumulation of 5’UTR, CDS and 3’UTR regions of *PYL6* mRNA compared between *xrn4-5* mutant and WT in response to ABA. This analysis represents three independent experiments. In panels B and C, seedlings were treated with 1 μM ABA for 1 hour before sampling. Values are the means of five biological replicates ± standard deviation. The relative levels of transcripts were obtained in comparison to the untreated WT. Responses were significantly different for fold changes in mRNA levels ≥ |1.5| and p < 0.05, according to Student’s t-test (* < 0.05; ** <0.005; *** < 0.0005).

First, we evaluated *PYL6* expression in the *dcp5-1* mutant, which is defective in a component of decapping machinery (Xu and Chua, 2009). *PYL5* was used as a control since it has been shown to be upregulated in *dcp5-1* (Wawer et al., 2018) and, indeed, *PYL5* transcript accumulated more in this mutant as compared to WT after ABA treatment (Fig. 4B). We found that ABA treatment resulted in 4.6-fold increase of *PYL6* mRNA levels in comparison to the WT, which suggests that the decapping machinery is involved in the downregulation of *PYL6* mRNA (Fig. 4B). In addition, *PYL4*, which belong to the same clade as *PYL5* and *PYL6* (Supplemental Fig. S1A), was also less repressed by ABA treatment in *dcp5-1* than in the WT (Fig. 4B). Thus, it is possible that ABA-promoted decapping may be a conserved mechanism for controlling the mRNA stability of these evolutionary related receptors.

To evaluated XRN4 involvement in the degradation of RISC 5’-fragment of *PYL6* mRNA, we carried out the quantification of the transcript level corresponding to different parts of *PYL6* transcript (5’UTR, CDS and 3’UTR) in the mutant *xrn4-5* in response to ABA. The amounts of all parts of *PYL6* mRNA were significantly more abundant in the *xrn4-5* mutant in comparison to WT (Fig. 4C). *PYL4* and *PYL5* transcripts were not altered in the mutant *xrn4-5* compared to WT (Supplemental Fig. S16). Since both 5’UTR and CDS regions from *PYL6* transcript are less efficiently reduced in *xrn4-5*,it is possible that XRN4 also participates in the degradation of *PYL6* mRNA from the 5’-end after decapping activity in response to ABA (Fig. 4C). Together, these results suggest 5’ to 3’ mRNA decay pathway is also involved in the degradation of *PYL6* transcript in response to ABA.

## DISCUSSION

Negative feedback is a key regulatory scheme in homeostatic processes such as those involved in hormone signaling (Teale et al., 2006; Zhu et al., 2013; Rai et al., 2015; Waters et al., 2017). After the initial increase in hormone levels in response to specific endogenous or exogenous signals, the hormone is detected by receptors and triggers adaptive responses through signaling pathways. These pathways need to be reset to maintain the hormone signaling homeostasis. In response to ABA, the negative feedback is achieved by down- and up-regulation of *PYR/PYL/RCAR* (positive regulators) and *PP2C* (negative regulators) gene expression, respectively (Song et al., 2016) (Fig 1A; Supplemental Fig. S1-S2). Our data highlight the requirement of a functional ABA signaling pathway for efficient ABA-induced downregulation of most *PYR/PYL/RCAR*receptors (Fig. 1C and 1D; Supplemental Fig. S4, S5 and S6). We further show that the control of mRNA decay is an essential step in shaping the ABA-induced repression of *PYL1* (subgroup III) and *PYL4/5/6* (subgroup II) expression (Fig. 1E; Supplemental Fig. S1A and Supplemental Table S3). These receptors have a lower affinity for ABA and were reported to be more active under abiotic stress conditions, when ABA levels are markedly increased (Tischer et al., 2017; Yoshida et al., 2019). Thus, ABA-induced degradation of *PYL1/4/5/6* mRNAs may be part of a strategy to avoid excessive and detrimental ABA responses under stress conditions.

The ABA-promoted decay processes of *PYL6* and *PYL1/4/5* transcripts are partly different from each other, since efficient repression of *PYL6* mRNA requires transcription to occur (Supplemental Table S3). This observation suggests that transcription of one causal agent involved in *PYL6* transcript destabilization is induced by the hormone (Fig. 1E; Supplemental Table S3). Therefore, we tested whether the miRNA pathway would be involved in *PYL6* mRNA destabilization in response to ABA. This hypothesis was supported by the impaired repression of *PYL6* mRNA levels by ABA in miRNA pathway mutants (Supplemental Fig. S8). MiR8175, miR5628, miR5021 and miR840-3p were identified as potential miRNAs acting upon *PYL6* transcript (Fig. 2A). Among these, several pieces of evidence indicate that miR5628 is involved in *PYL6* mRNA decay in response to ABA. First, as expected, both *pri-miR5628* and *miR5628* were quickly and transiently induced by ABA (Fig. 2D). This induction coincides with the fast reduction of *PYL6* transcript after ABA addition (2-fold of reduction in 10 minutes; Fig. 4A) and requires functional ABA signaling as revealed by the reduced accumulation of miR5628 in *snrk2d* mutant (Fig. 2E). Second, a fusion transcript consisting of GFP and *PYL6* coding sequence, but lacking miRNA5628 target in *PYL6* 3’UTR, was not repressed by ABA (Fig. 2B). Third, the transient expression assay in *N. benthamiana* of a luciferase construct incorporating miR5628 target site in the 3’UTR of LUC gene shown a tendency to reduce the luciferase activity when co-expressed with miR5628 gene (Supplemental Fig. S10D). Fourth, *PYL6* was more repressed during the first 20 minutes of ABA treatment in the miR5628 overexpressing lines (oxMIR5628-E4 and oxMIR5628-E6) compared to WT (Fig 2C).In addition, 5’-RACE products obtained from WT and these over-expressor lines were found to map to the miR5628 recognition site in *PYL6* transcript or downstream to it, but none was found to map upstream (Fig. 2A and Supplemental Fig. S12). Fifth, germination of oxMIR5628 seeds was found to be hyposensitive to ABA (Fig 3A). Thus, it is suggested that miR5628-mediating *PYL6* repression impacts ABA sensitivity at this developmental stage. Finally, XRN4, which is known to degrade RISC 3’-cleaved fragments of miRNA-targets (Souret et al., 2004), was found to be required for efficient ABA-induced decay of *PYL6*-3’UTR mRNA region (Supplemental Fig. S10B). Together, these set of observations support the notion that miR5628 promotes *PYL6* mRNA degradation by promoting its cleavage in an ABA-dependent manner (Fig. 5).

**Figure 5:**
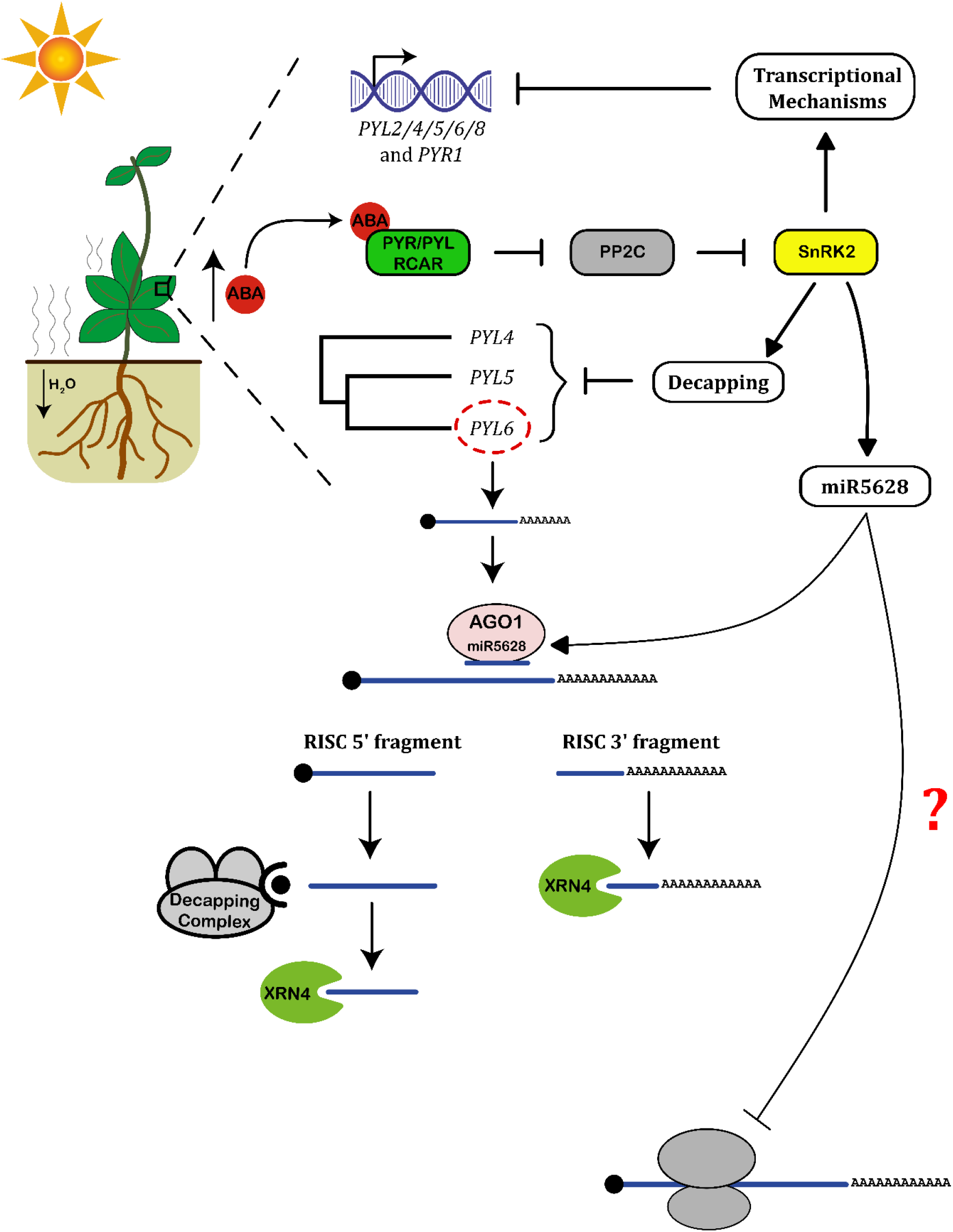
Model of the control of ABA signalization through repression of *PYR/PYL/RCAR* genes. Abiotic stress conditions such as drought, increase the endogenous level of ABA, which is perceived by PYR/PYL/RCAR receptors. The ABA-receptor complex sequesters Clade A PP2C phosphatases, releasing SnRK2 kinases from their negative regulation. These SnRK2s phosphorylate several proteins in order to activate the gene expression program of ABA responses. Part of these ABA responses are involved in the transcriptional repression of *PYL1/2/4/5/6* and *PYL8* genes. In additicn, ABA accelerates the decay of *PYL1/4/5/6* transcripts. The ABA core signaling pathway induces miR5628 expression, which in turn is processed and loaded onto AG01. AG01-miR5628 complex promotes the cleavage of *PYL6* mRNA at the 3’UTR region, and XRN4 promotes the degradation of the RISC 3’-cleaved fragment of *PYL6*. Additionally, the dynamic of *PYL6* mRNA decay may involve the participation of 5’ to 3’ mRNA decay pathway, where RISC 5’-cleaved fragment of *PYL6* transcripts would undergo decapping followed by XRN4-mediate degradation. Decapping may also contribute to destabilize *PYL4* and *PYL5* transcripts, which are phylogenetically close to *PYL6*. The repression of *PYR/PYL/RCAR* genes expression is a mean to limit de novo synthesis of receptors, controlling the extension of ABA responses and participating in the resetting of the ABA signaling. In addition, it might be possible that miR5628 would regulate *PYL6* mRNA translation.

MiR5628 is likely to be restricted to *A. thaliana* and is predicted to specifically recognizes *PYL6* mRNA among the 14 *PYR/PYL/RCAR* members (Supplemental Fig. S14 and S15). Therefore, we hypothesized that miR5628 is a regulatory novelty that was integrated into the feedback loop of ABA signaling to enhance *PYL6* mRNA decay. This hypothesis would possibly explain the faster and stronger ABA-promoted *PYL6* mRNA decay compared to the others *PYR/PYL/RCAR* transcripts (Fig. 1E, Supplemental Fig. S7 and Supplemental Table S3).

Intriguingly, miR5628-guided cleavage at the expected canonical position (*i.e*., 10 and 11^th^ nucleotide) was underrepresented in the 5’ RACE clones (Supplemental Fig. S12). Such features have been described for other miRNAs (Fahlgren et al., 2007; Lee et al. 2015; Gou et al., 2022; Ren et al., 2022) and, although the underlying reasons are unclear, they would be partly related to peculiarities of newly evolved miRNA (Fahlgren et al., 2012; Axtell, 2013; Hajieghrari and Farrokhi, 2021). In the case of miR5628, some features such as incomplete pairing with its target (Supplemental Fig. S9) and the suboptimal guanine at its 5’-end for AG01 loading which would affect the stability of miRNA-target duplex, could be responsible for reducing its cleavage activity and contribute to the looser definition of target site cutting (Mi et al., 2008). In addition, it is also possible that the putative secondary structure of *PYL6* mRNA can modulate its recognition by miR5628 (Supplemental Fig. S17)(Zheng et al., 2017).

The ABA signaling readouts genes *RD29B, RAB18* and *RD20* were upregulated in the *oxPYL6* genotype either in the absence or presence of ABA (Fig. 3B), most likely as a consequence of more receptors triggering ABA signaling. In the oxMIR5628 lines, ABA does not affect the expression of these readout genes as compared to the WT (Figure 3C), possibly as a consequence of functional redundancy among PYL/PYR/RCAR receptors (Zhao et al., 2018). In the absence of ABA, these readout genes were downregulated in oxMIR5628 lines (Fig. 3C), yet *PYL6* mRNA levels was not affected (Fig. 2C). The reason for this inconsistency is not clear but may be related to the possibility of miR5628-regulating *PYL6* mRNA translation, since it has been reported that miRNAs which targets 3’-UTR regions, as is the case for miR5628, inhibit the translation of their mRNA targets (Gandikota et al., 2007; Iwakawa and Tomari, 2013).

Following miR5628-mediated cleavage of *PYL6* mRNA in response to ABA, we expected that a faster decay of the *PYL6*-3’UTR region in comparison to the 5’UTR and CDS in response to ABA would occur. However, a synchronized pattern of degradation of the 5’UTR, CDS and 3’UTR regions of *PYL6* mRNA after ABA treatment was observed (Fig. 4A), suggesting that, in addition to miR5628 activity, other post-transcriptional regulatory mechanisms would participate in the regulation of *PYL6* transcript decay.

Indeed, we provided evidence that the decapping machinery is involved in the ABA-induced reduction of *PYL6* mRNA levels (Fig. 4B). In addition to the degradation of RISC 3’-cleaved fragment of *PYL6*-3’UTR, XRN4 is also involved in the degradation of *PYL6* 5’UTR and CDS regions (Fig. 4C). These data raise the hypothesis that miR5628-guided *PYL6* mRNA cleavage together with decapping and XRN4 activity would be coupled processes that promote *PYL6* transcript degradation in response to ABA (Fig. 5).

Similarly to *PYL6* transcript, the control of mRNA decay of *PYL4/5* receptors, at least in part, rely on decapping activity, since these transcripts were shown to be less responsive to ABA in the decapping *dcp5-1* mutant (Fig. 4B). This result is in line with the observation that the mRNA of these two receptors and of *PYL6* are targets of VARICOSE, another component of decapping machinery (Sorenson et al., 2018). Since *PYL4/5/6* are part of the same clade in subgroup III (Supplemental Fig. S1A), we suggest that the involvement of decapping in the control of the stability of these ABA receptors transcripts is an ancestral regulatory feature.

The control of PYR/PYL/RCAR protein stability is another aspect of the dynamic of regulation of the ABA signaling. Most PYR/PYL/RCAR receptors tend to be ubiquitinated and degraded by the 26S proteasome in the absence of ABA and stabilized in the presence of ABA (Irigoyen et al., 2014; Chen et al., 2018; Li et al., 2018). This regulation favors the establishment of ABA-induced gene expression programs. Fast ABA-induced repression of *PYL1, PYL4, PYL5* and *PYL6* transcripts levels through, at least in part, the control of mRNA stability, would counterbalance receptors stabilization and contribute to limit *de novo* synthesis of receptors and, thus, constrain ABA-responses. Hence, the balance between PYR/PYL/RCAR protein stability and the level of the corresponding mRNA would define the extent of ABA signalization and is likely an important facet in the homeostasis of ABA signaling.

## MATERIAL AND METHODS

### Plant material and growth conditions

Five milligrams of seeds were surface-sterilized and added to 10 mL of half-strength liquid Murashige and Skoog medium (MS/2) adjusted to 0.3% of Glucose (Glc) (w/v). After stratification at 4°C for three days, seedlings were grown under constant light (Photosynthetically Active Radiation/PAR of 100 μmol m^-2^s^-1^) at 21°C for 6 days under constant agitation (60 rpm). On the sixth day, samples were treated with 1 μM ABA (final concentration) for different times. Five biological replicates per treatment were used. The following mutant lines in the Col-0 background were used: *abi1-1* (Umezawa et al., 2009), *abi4-1* (Finkelstein, 1994), *snrk2d* (Fujii et al., 2007) *se-1, hyl1-2* (Vazquez et al., 2004), *ago1-25* (Morel et al., 2002), *dcp5-1* (Xu and Chua, 2009) and *xrn4-5* (Souret et al., 2004). Seeds of Col-0 background containing the fusion between the coding sequence of the Green Fluorescent Protein (GFP) and of PYL6 under the control of the *Cauliflower Mosaic Virus (CaMV*) 35S promoter (*oxPYL6*) were described in Belda-Palazon et al. (2016).

*Nicotiana benthamiana* plants were grown at 22°C under relative humidity 65% and 16h/8h (light/dark) conditions on plastic cups containing peat:vermiculite (1:1). Plants at six weeks after germination were used for Agroinfiltration.

### Seed germination analysis

Surface-sterilized seeds were sawn on half-strength solid MS adjusted to 0.3% Glc (w/v) and grown under continuous light (PAR of 100 μmol m^-2^s^-1^). Seed germination rate (%) corresponds to the number of seeds that germinated each day for the total of seeds germinated at the sixth day. We apply Tukey and Scott-Knott test to evaluate significant differences in the germination rate.

### RNA isolation and gene expression analysis

Total RNA extraction, cDNA synthesis and real-time quantitative reverse transcription polymerase chain reaction (RT-qPCR) were conducted as previously described (Matiolli et al., 2011; Duarte et al., 2013). Relative quantification levels were calculated by the 2 ^-ΔΔCT^ formula. To quantify different parts of *PYL6* and *GFP* transcripts in WT and oxPYL6 genotypes the formula 2^(Ct target gene - Ct reference genes)*100^ was used. The reference genes were *PDF2* (AT1G13320) and *EF-1α* (AT5G60390) (Czechowski et al., 2005). Stem-loop RT-qPCR miRNA assay was carried out for the quantification of mature miRNA (Varkonyi-Gasic et al., 2007). Differences in gene expression were considered significant for fold changes ≥ |1.5| between treated and control samples and for p < 0.05 according to two-tailed Student’s t test. The primer sequences for RT-qPCR are shown in Supplemental Table S1.

### Staurosporine and Cordycepin treatments

Five days-old seedlings were treated for 24 h with 10 μM staurosporine (stock solution 100 mM in dimethylsulfoxide; Sigma S5921). On the sixth day, the seedlings were treated with 1 μM ABA for 1 hour (h). Control samples were treated with DMSO.

Transcription inhibition was performed with 100 μM cordycepin (3-deoxyadenosine; Sigma C3394) from a 100 mM stock aqueous solution. To evaluate ABA-mediated control of mRNA stability we proceed according describe by Matiolli et al. (2011).

### 5’-RACE analysis

5’RACE was performed using the GeneRacer kit (Invitrogen) according to the manufacturer’s recommendations. Five micrograms of total RNA were ligated to the RNA GeneRacer oligo adapter and subjected to reverse transcription utilizing Improm-II Reverse Transcriptase kit (Promega). The cDNA was used for amplification of cleaved *PYL6* fragments, using a forward primer specific for the sequence of the GeneRacer RNA oligo adapter and a reverse primer specific for the *PYL6* mRNA (Supplemental Table S2). PCR products were then used as a template for a NESTED PCR with internal *PYL6* specific primers (Supplemental Table S2). After amplification, 5’RACE products were gel-purified and cloned in pGem®-T Easy vector (Promega). Independent clones were randomly chosen and sequenced by Sanger sequencing method.

### DNA constructions

Plasmids used in this study were constructed by modifying the pGreen dual-Luc 3’-UTR sensor plasmid by double digested with Agel and AvrII restriction enzymes, as early described (Liu et al., 2014). T4 DNA ligase-mediated insertion of *PYL6* target sequence (WT sequence), miR5628 perfect match (positive control or non-complementary sequence of miR5628 (negative control) into the 3’-UTR pGreen dual-Luc plasmid was performed and Sanger sequenced. Versions of miR5628 targets sites were obtained by annealing two complementary primers (Supplemental Table S2). For precursor of miR5628 of *A. thaliana*, we designed forward and reverse primers 180 and 122 nucleotides up and downstream to the precursor sequence of miR5628, respectively (Supplemental Table S2). Products of PCR amplified were sub-cloned into pENTR™ Directional TOPO^®^ and further Sanger sequenced followed by cloning into the Gateway pK7WG2.0.

### Agroinfiltration and Dual-luciferase assay

Plasmids were introduced by electroporation into *Agrobacterium tumefaciens* strain GV3101 (harboring pSOUP) and plated on LB agar broth containing rifampicin (25 μg/mL), gentamicin (25 μg/mL) and kanamycin (50 μg/mL) selection. Primary inoculations were prepared by inoculating a single colony and grown overnight at 28°C in a shaking incubator. Working cultures were harvested by centrifugation at room temperature at 3000 rpm for 10 min. Cell pellets were resuspended in 2 mL of room temperature infiltration media (88.5 mL water, 1 mL 1M MgCl2, 10 mL 100 mM MES, 75 μL 200 mM acetosyringone) and stored in the dark at room temperature for 4 h. The OD_600_ were adjusted to about 0.5. In three 15-mL falcon tubes it was mixed OD_600_-adjusted sensor and p35S::MIR5628 culture per tube at 1:1 ratio. Three expanded leaves per plant were infiltrated by applying pressure on the abaxial surface of the leaf with a 1-mL syringe with well-mixed *Agrobacterium* suspension.

After 3 days of agroinfiltration treatment it was punched two leaf discs from each leaf and placed into 1.5 mL eppendorf tube. Samples were frozen immediately into liquid nitrogen and grinded. Fine power was resuspended into the ice-chilled lysis buffer (PLB) and shacked on vortex to completely resuspend the tissue powder in the solution. Resuspended samples were centrifuged at 14000 rpm, 4°C, for 1 min to pellet cell debris. It was loaded 20 μL of the supernatant from each sample into designated position on the 96-well plate. The substrate solution LARII was prepared for firefly Luciferase, and the substrate solution Stop&Glo was used for Renilla. The analysis was performed on GloMax 96 Microplate Luminometer (Promega), and the F-Luc/R-Luc ratio was calculated for all samples and technical replicates.

### Generation of Transgenic lines

To make oxMIR5628 lines, the vector pK7WG2.0, containing the 35S:MIR5628 fusion, were introduced by electroporation into *A. tumefaciens* strain GV3101 and plated on LB agar broth containing rifampicin (25 μg/mL) and kanamycin (50 μg/mL) selection. Then, Arabidopsis plants (Col-0 background) were transformed according floral dip method. Transformants were selected by their kanamycin (50 μg/mL) resistance and validated by PCR. Homozygous lines (oxMIR5628) were used for experiments.

### Bioinformatics analysis

The software psRNA-target was used to identify putative miRNAs that can recognize *PYL6* transcript (http://plantnrn.noble.org/psRNATarnet/home) (Dai et al., 2018). Analysis of miRNA conservation was carried out using BLAST. Both precursor (pre-miR5628) and mature miR5628 sequences were used as query to search for homologs in the genomes of the vegetal kingdom available at the National Center for Biotechnology Information (NCBI) (https://www.ncbi.nlm.nih.gov/). Blast parameters were adjusted as follows: expect values were set at 1000; high similar sequences were chosen as the sequence filter; the number of descriptions and alignments was raised to 1000 and the default word-match size between the query and database sequences was seven. The ability of sequences similar to pre-miR5628 to form stem-loop structure *in silico* was evaluated using the RNAfold web server (http://rna.tbi.univie.ac.at/) (Gruber et al., 2008) and their expression was verified searching transcriptomes data available in the Sequence Read Archive (RSA) at the NCBI and. We analyzed the global miR5628 expression available in the *Arabidopsis Small RNA Database* (http://ipf.sustech.edu.cn/pub/asrd/) (Feng et al., 2020).

Amino acid sequences of the 14 *A. thaliana* PYR/PYL/RCAR were retrieved from the TAIR10 database (https://www.arabidopsis.org/). These sequences were used to infer the phylogenetic relationship of these proteins using Neighbor-Joining method and draw the corresponding tree of similarity using MEGA7 program (Kumar et al., 2016).

## ACKNOWLEDGEMENTS

The authors are grateful to Hervé Vaucheret, Baena-Gonzáles, Jian-Kang Zhu and Pedro Rodriguez for providing seeds of the different genotypes used in this work. This work was funded by Fundação de Amparo à Pesquisa do Estado de São Paulo (FAPESP): Vincentz, M (grant 2015/25838-2), Vieira JGP (grant 2016/0498-7 and 2019/25696-4) and Duarte, GT (grant 12/22125-7).

## SUPPLEMENTAL DATA

**Supplemental Table S1:** Sequence of oligonucleotides used for RT-qPCR analysis. **Supplemental Table S2:**Sequence of oligonucleotides used for 5’RACE and Dual-luciferase analyses.

**Supplemental Table S3:** ABA-induced changes in the mRNA stability of PYR/PYL/RCAR.

**Supplemental Table S4:** Global expression of miR5628 in comparison to newly evolved and conserved miRNAs in *Arabidopsis thaliana*.

**Supplemental Figure S1:** Effect of ABA treatment on gene expression of ABA core signaling components.

**Supplemental Figure S2:** mRNA profiles of ABA core signaling pathway genes in response to long-term treatment with ABA.

**Supplemental Figure S3:** Changes in the expression profile of ABA core signaling pathway genes in response to a short ABA treatment.

**Supplemental Figure S4:** Downregulation of *PYR/PYL/RCAR* gene expression requires a functional ABA core signaling pathway.

**Supplemental Figure S5:** Global kinase inhibition by staurosporine affects the expression of *PYR/PYL/RCAR* and clade A *PP2C* genes in response to ABA.

**Supplemental Figure S6:** Kinases SnRK2 from subclass III are involved in the downregulation of *PYR/PYL/RCAR* gene expression in response to ABA.

**Supplemental Figure S7:** Impact of ABA on the *PYR1, PYL2* and *PYL8* mRNA stability.

**Supplemental Figure S8:** Involvement of miRNA pathway in the control of *PYL6* expression in response to ABA.

**Supplemental Figure S9:** Schematic representation of *PYL6* mRNA and the position of the putative miR8175, miR5628, miR5021 and miR840-3p target sequences.

**Supplemental Figure S10:** Analyses of cleavage of *PYL6* mRNA by miR5628.

**Supplemental Figure S11:** Lineages oxMIR5628-E4 and oxMIR5628-E6 overexpressed both the primary miR5628 sequence (*pri-miR5628*) and mature miR5628 sequence (*miR5628*).

**Supplemental Figure S12:** Schematic representation of 5’RACE cloned sequences of *PYL6* mRNA upon 20 minutes of ABA treatment in the oxMIR5628 (E4 and E6) lineages and Col-0.

**Supplemental Figure S13:** MiR5628 biogenesis is controlled by ABA.

**Supplemental Figure S14:** Divergence of the miR5628 recognition sequence among the 14 *A. thaliana* ABA receptors.

**Supplemental Figure S15:** Overexpression of miR5628 do not impact the expression of *PYR1, PYL1, PYL2, PYL4, PYL5* and *PYL8*.

**Supplemental Figure S16:** Analysis of *PYL4* and *PYL5* mRNA accumulation in wild type (WT) and *xrn4-5* in response to ABA treatment.

**Supplemental Figure S17:** Predicted *PYL6* mRNA 3’-UTR secondary structure includes miR5628 target site.

## Parsed Citations

**Axtell MJ (2013) Classification and comparison of small RNAs from plants. Annu Rev Plant Biol 64: 137–159**

Google Scholar: <u>Author Only Title Only Author and Title</u>

**Barrera-Rojas CH, Otoni WC, Nogueira FTS (2021) Shaping the root system: the interplay between miRNA regulatory hubs and phytohormones. J Exp Bot 72: 6822–6835**

Google Scholar: <u>Author Only Title Only Author and Title</u>

**Belda-Palazon B, Rodriguez L, Fernandez MA, Castillo M-C, Anderson EA, Gao C, González-Guzmán M, Peirats-Llobet M, Zhao Q, De Winne N, et al (2016) FYVE1/FREE1 Interacts with the PYL4 ABA Receptor and Mediates its Delivery to the Vacuolar Degradation Pathway. Plant Cell 28: tpc.00178.2016**

Google Scholar: <u>Author Only Title Only Author and Title</u>

**Chantarachot T, Bailey-Serres J (2018) Polysomes, Stress Granules, and Processing Bodies: A Dynamic Triumvirate Controlling Cytoplasmic mRNA Fate and Function. Plant Physiol 176: 254–269**

Google Scholar: Author Only Title Only Author and Title

**Chen H-H, Qu L, Xu Z-H, Zhu J-K, Xue H-W (2018) EL1-like Casein Kinases Suppress ABA Signaling and Responses by Phosphorylating and Destabilizing the ABA Receptors PYR/PYLs in Arabidopsis. Mol Plant 11: 706–719**

Google Scholar: <u>Author Only Title Only Author and Title</u>

**Chu LH, Jeng ST (2002) Multiple transduction pathways regulate the 35S promoter with an ABA responsive element. Plant Sci 163: 23–32**

Google Scholar: <u>Author Only Title Only Author and Title</u>

**Czechowski T, Stitt M, Altmann T, Udvardi MK, Scheible W-R (2005) Genome-wide identification and testing of superior reference genes for transcript normalization in Arabidopsis. Plant Physiol 139: 5–17**

Google Scholar: <u>Author Only Title Only Author and Title</u>

**Dai X, Zhuang Z, Zhao PX (2018) psRNATarget: a plant small RNA target analysis server (2017 release). Nucleic Acids Res 46: W49–W54**

Google Scholar: <u>Author Only Title Only Author and Title</u>

**Davière J-M, Achard P (2013) Gibberellin signaling in plants. Development 140: 1147–1151**

Google Scholar: <u>Author Only Title Only Author and Title</u>

**Duarte GT, Matiolli CC, Pant BD, Schlereth A, Scheible W-RW-R, Stitt M, Vicentini R, Vincentz M (2013) Involvement of microRNA-related regulatory pathways in the glucose-mediated control of Arabidopsis early seedling development. J Exp Bot 64: 4301–4312**

Google Scholar: <u>Author Only Title Only Author and Title</u>

**Fahlgren N, Howell MD, Kasschau KD, Chapman EJ, Sullivan CM, Cumbie JS, Givan SA, Law TF, Grant SR, Dangl JL, et al (2007) High-Throughput Sequencing of Arabidopsis microRNAs: Evidence for Frequent Birth and Death of MIRNA Genes. PLoS One 2: e219**

Google Scholar: <u>Author Only Title Only Author and Title</u>

**Fahlgren N, Jogdeo S, Kasschau KD, Sullivan CM, Chapman EJ, Laubinger S, Smith LM, Dasenko M, Givan SA, Weigel D, et al (2012) MicroRNA Gene Evolution in Arabidopsis lyrata and Arabidopsis thaliana. Plant Cell 22: 1074–1089**

Google Scholar: <u>Author Only Title Only Author and Title</u>

**Feng L, Zhang F, Zhang H, Zhao Y, Meyers BC, Zhai J (2020) An Online Database for Exploring Over 2,000 Arabidopsis Small RNA Libraries. Plant Physiol 182: 685–691**

Google Scholar: <u>Author Only Title Only Author and Title</u>

**Finkelstein RR (1994) Mutations at two new Arabidopsis ABA response loci are similar to the abi3 mutations. Plant J 5: 765–771**

Google Scholar: <u>Author Only Title Only Author and Title</u>

**Fujii H, Verslues PE, Zhu JK (2007) Identification of Two Protein Kinases Required for Abscisic Acid Regulation of Seed Germination, Root Growth, and Gene Expression in Arabidopsis. Plant Cell 19: 485–494**

Google Scholar: <u>Author Only Title Only Author and Title</u>

**Fujita Y, Nakashima K, Yoshida T, Katagiri T, Kidokoro S, Kanamori N, Umezawa T, Fujita M, Maruyama K, Ishiyama K, et al (2009) Three SnRK2 Protein Kinases are the Main Positive Regulators of Abscisic Acid Signaling in Response to Water Stress in Arabidopsis. Plant Cell Physiol 50: 2123–2132**

Google Scholar: <u>Author Only Title Only Author and Title</u>

**Gandikota M, Birkenbihl RP, Höhmann S, Cardon GH, Saedler H, Huijser P (2007) The miRNA156/157 recognition element in the 3’ UTR of the Arabidopsis SBP box gene SPL3 prevents early flowering by translational inhibition in seedlings. Plant J 49: 683–693**

Google Scholar: <u>Author Only Title Only Author and Title</u>

**Gou X, Zhong C, Zhang P, Mi L, Li Y, Lu W, Zheng J, Xu J, Meng Y, Shan W (2022) miR398b and AtC2GnT form a negative feedback loop to regulate Arabidopsis thaliana resistance against Phytophthora parasitica. Plant J 111: 360–373**

Google Scholar: <u>Author Only Title Only Author and Title</u>

**Gruber AR, Lorenz R, Bernhart SH, Neubock R, Hofacker IL (2008) The Vienna RNA Websuite. Nucleic Acids Res 36: W70–W74**

Google Scholar: <u>Author Only Title Only Author and Title</u>

**Hajieghrari B, Farrokhi N (2021) Investigation on the Conserved MicroRNA Genes in Higher Plants. Plant Mol Biol Report 39: 10– 23**

Google Scholar: <u>Author Only Title Only Author and Title</u>

**Irigoyen ML, Iniesto E, Rodriguez L, Puga MI, Yanagawa Y, Pick E, Strickland E, Paz-Ares J, Wei N, De Jaeger G, et al (2014) Targeted degradation of abscisic acid receptors is mediated by the ubiquitin ligase substrate adaptor DDA1 in Arabidopsis. Plant Cell 26: 712–28**

Google Scholar: <u>Author Only Title Only Author and Title</u>

**Iwakawa H oki, Tomari Y (2013) Molecular Insights into microRNA-Mediated Translational Repression in Plants. Mol Cell 52: 591– 601**

Google Scholar: <u>Author Only Title Only Author and Title</u>

**Kavi Kishor PB, Tiozon RN, Fernie AR, Sreenivasulu N (2022) Abscisic acid and its role in the modulation of plant growth, development, and yield stability. Trends Plant Sci 27: 1283–1295**

Google Scholar: <u>Author Only Title Only Author and Title</u>

**Kieber JJ, Schaller GE (2018) Cytokinin signaling in plant development. Development 145: dev149344**

Google Scholar: <u>Author Only Title Only Author and Title</u>

**Klepikova A V., Kasianov AS, Gerasimov ES, Logacheva MD, Penin AA (2016) A high resolution map of the Arabidopsis thaliana developmental transcriptome based on RNA-seq profiling. Plant J 88: 1058–1070**

Google Scholar: <u>Author Only Title Only Author and Title</u>

**Kumar S, Stecher G, Tamura K (2016) MEGA7: Molecular Evolutionary Genetics Analysis Version 7.0 for Bigger Datasets. Mol Biol Evol 33: 1870–1874**

Google Scholar: <u>Author Only Title Only Author and Title</u>

**Lee HJ, Park YJ, Kwak KJ, Kim D, Park JH, Lim JY, Shin C, Yang KY, Kang H (2015) MicroRNA844-guided downregulation of cytidinephosphate diacylglycerol Synthase3 (CDS3) mRNA affects the response of arabidopsis thaliana to bacteria and fungi. Mol Plant-Microbe Interact 28: 892–900**

Google Scholar: <u>Author Only Title Only Author and Title</u>

**Li D, Zhang L, Li X, Kong X, Wang X, Li Y, Liu Z, Wang J, Li X, Yang Y (2018) AtRAE1 is involved in degradation of ABA receptor RCAR1 and negatively regulates ABA signalling in Arabidopsis. Plant Cell Environ 41: 231–244**

Google Scholar: <u>Author Only Title Only Author and Title</u>

**Liu Q, Wang F, Axtell MJ (2014) Analysis of complementarity requirements for plant microRNA targeting using a Nicotiana benthamiana quantitative transient assay. Plant Cell 26: 741–53**

Google Scholar: <u>Author Only Title Only Author and Title</u>

**Ma Y, Szostkiewicz I, Korte A, Moes D, Yang Y, Christmann A, Grill E (2009) Regulators of PP2C Phosphatase Activity Function as Abscisic Acid Sensors. Science (80-) 324: 1064–1069**

Google Scholar: <u>Author Only Title Only Author and Title</u>

**Matiolli CC, Tomaz JP, Duarte GT, Prado FM, Del Bem LEV, Silveira AB, Gauer L, Corrêa LGG, Drumond RD, Viana AJC, et al (2011) The Arabidopsis bZIP gene AtbZIP63 is a sensitive integrator of transient abscisic acid and glucose signals. Plant Physiol 157: 692–705**

Google Scholar: <u>Author Only Title Only Author and Title</u>

**Mi S, Cai T, Hu Y, Chen Y, Hodges E, Ni F, Wu L, Li S, Zhou H, Long C, et al (2008) Sorting of small RNAs into Arabidopsis argonaute complexes is directed by the 5’ terminal nucleotide. Cell 133: 116–127**

Google Scholar: <u>Author Only Title Only Author and Title</u>

**Morel J-BJ-B, Godon C, Mourrain P, Béclin C, Boutet S, Feuerbach F, Proux F, Vaucheret H (2002) Fertile hypomorphic ARGONAUTE (ago1) mutants impaired in post-transcriptional gene silencing and virus resistance. Plant Cell 14: 629–39**

Google Scholar: <u>Author Only Title Only Author and Title</u>

**Mugridge JS, Coller J, Gross JD (2018) Structural and molecular mechanisms for the control of eukaryotic 5’–3’ mRNA decay. Nat Struct Mol Biol 25: 1077–1085**

Google Scholar: <u>Author Only Title Only Author and Title</u>

**Park S, Fung P, Nishimura N, Jensen DR, Fujii H, Zhao Y, Lumba S, Santiago J, Rodrigues A, Chow TF, et al (2009) Abscisic Acid**

**Inhibits Type 2C Protein Phosphatases via the PYR/PYL Family of START Proteins. Science (80-) 324: 1068–1069**

Google Scholar: <u>Author Only Title Only Author and Title</u>

**Rai MI, Wang X, Thibault DM, Kim HJ, Bombyk MM, Binder BM, Shakeel SN, Schaller GE (2015) The ARGOS gene family functions in a negative feedback loop to desensitize plants to ethylene. BMC Plant Biol 15: 157**

Google Scholar: <u>Author Only Title Only Author and Title</u>

**Ren G, Meng X, Shuxin Z, Carissa V, Chen X, Bin. Y (2014) Methylation protects microRNAs from an AGO1-associated activity that uridylates 5’ RNA fragments generated by AGO1 cleavage. Proc Natl Acad Sci 111: 6365–6370**

Google Scholar: <u>Author Only Title Only Author and Title</u>

**Ren Y, Li M, Wang W, Lan W, Schenke D, Cai D, Miao Y (2022) MicroRNA840 (MIR840) accelerates leaf senescence by targeting the overlapping 3’UTRs of PPR and WHIRLY3 in Arabidopsis thaliana. Plant J 109: 126–143**

Google Scholar: <u>Author Only Title Only Author and Title</u>

**Somvanshi PR, Patel AK, Bhartiya S, Venkatesh K V. (2015) Implementation of integral feedback control in biological systems. Wiley Interdiscip Rev Syst Biol Med 7: 301–316**

Google Scholar: <u>Author Only Title Only Author and Title</u>

**Song L, Huang SC, Wise A, Castanon R, Nery JR, Chen H, Watanabe M, Thomas J, Bar-Joseph Z, Ecker JR (2016) A transcription factor hierarchy defines an environmental stress response network. Science. doi: 10.1126/science.aag1550**

Google Scholar: <u>Author Only Title Only Author and Title</u>

**Sorenson RS, Deshotel MJ, Johnson K, Adler FR, Sieburth LE (2018) Arabidopsis mRNA decay landscape arises from specialized RNA decay substrates, decapping-mediated feedback, and redundancy. Proc Natl Acad Sci U S A 115: E1485–E1494**

Google Scholar: <u>Author Only Title Only Author and Title</u>

**Souret FF, Kastenmayer JP, Green PJ (2004) AtXRN4 Degrades mRNA in Arabidopsis and Its Substrates Include Selected miRNA Targets. Mol Cell 15: 173–183**

Google Scholar: <u>Author Only Title Only Author and Title</u>

**Teale WD, Paponov IA, Palme K (2006) Auxin in action: signalling, transport and the control of plant growth and development. Nat Rev Mol Cell Biol 7: 847–859**

Google Scholar: <u>Author Only Title Only Author and Title</u>

**Tischer S V, Wunschel C, Papacek M, Kleigrewe K, Hofmann T, Christmann A, Grill E (2017) Combinatorial interaction network of abscisic acid receptors and coreceptors fromArabidopsis thaliana. Proc Natl Acad Sci U S A 114: 10280–10285**

Google Scholar: <u>Author Only Title Only Author and Title</u>

**Umezawa T, Sugiyama N, Mizoguchi M, Hayashi S, Myouga F, Yamaguchi-Shinozaki K, Ishihama Y, Hirayama T, Shinozaki K (2009) Type 2C protein phosphatases directly regulate abscisic acid-activated protein kinases in Arabidopsis. Proc Natl Acad Sci U S A 106: 17588–93**

Google Scholar: <u>Author Only Title Only Author and Title</u>

**Urano K, Maruyama K, Jikumaru Y, Kamiya Y, Yamaguchi-Shinozaki K, Shinozaki K (2017) Analysis of plant hormone profiles in response to moderate dehydration stress. Plant J 90: 17–36**

Google Scholar: <u>Author Only Title Only Author and Title</u>

**Varkonyi-Gasic E, Wu R, Wood M, Walton EF, Hellens RP (2007) Protocol: a highly sensitive RT-PCR method for detection and quantification of microRNAs. Plant Methods 3: 12**

Google Scholar: <u>Author Only Title Only Author and Title</u>

**Vazquez F, Gasciolli V, Crété P, Vaucheret H (2004) The Nuclear dsRNA Binding Protein HYL1 Is Required for MicroRNA Accumulation and Plant Development, but Not Posttranscriptional Transgene Silencing. Curr Biol. doi: 10.1016/j.cub.2004.01.035**

Google Scholar: <u>Author Only Title Only Author and Title</u>

**Wang J-W, Czech B, Weigel D (2009) miR156-Regulated SPL Transcription Factors Define an Endogenous Flowering Pathway in Arabidopsis thaliana. Cell 138: 738–749**

Google Scholar: <u>Author Only Title Only Author and Title</u>

**Wang P, Xue L, Batelli G, Lee S, Hou Y-J, Van Oosten MJ, Zhang H, Tao WA, Zhu J-K (2013) Quantitative phosphoproteomics identifies SnRK2 protein kinase substrates and reveals the effectors of abscisic acid action. Proc Natl Acad Sci U S A 110: 11205–10**

Google Scholar: <u>Author Only Title Only Author and Title</u>

**Wang Z, Ji H, Yuan B, Wang S, Su C, Yao B, Zhao H, Li X (2015) ABA signalling is fine-tuned by antagonistic HAB1 variants. Nat Commun 6: 8138**

Google Scholar: <u>Author Only Title Only Author and Title</u>

**Waters MT, Gutjahr C, Bennett T, Nelson DC (2017) Strigolactone Signaling and Evolution. Annu Rev Plant Biol 68: 291–322**

Google Scholar: <u>Author Only Title Only Author and Title</u>

**Wawer I, Golisz A, Sulkowska A, Kawa D, Kulik A, Kufel J (2018) mRNA decapping and 5’-3’ decay contribute to the regulation of ABA signaling in Arabidopsis thaliana. Front Plant Sci 9: 312**

Google Scholar: <u>Author Only Title Only Author and Title</u>

**Xu J, Chua N-H (2009) Arabidopsis decapping 5 is required for mRNA decapping, P-body formation, and translational repression during postembryonic development. Plant Cell 21: 3270–9**

Google Scholar: <u>Author Only Title Only Author and Title</u>

**Yoshida T, Christmann A, Yamaguchi-Shinozaki K, Grill E, Fernie AR (2019) Revisiting the Basal Role of ABA - Roles Outside of Stress. Trends Plant Sci. doi: 10.1016/j.tplants.2019.04.008**

Google Scholar: <u>Author Only Title Only Author and Title</u>

**Yoshida T, Fujita Y, Maruyama K, Mogami J, Todaka D, Shinozaki K, Yamaguchi-Shinozaki K (2015) Four Arabidopsis AREB/ABF transcription factors function predominantly in gene expression downstream of SnRK2 kinases in abscisic acid signalling in response to osmotic stress. Plant Cell Environ 38: 35–49**

Google Scholar: <u>Author Only Title Only Author and Title</u>

**Zhao Y, Zhang Z, Gao J, Wang P, Hu T, Wang Z, Hou YJ, Wan Y, Liu W, Xie S, et al (2018) Arabidopsis Duodecuple Mutant of PYL ABA Receptors Reveals PYL Repression of ABA-Independent SnRK2 Activity. Cell Rep 23: 3340-3351.e5**

Google Scholar: <u>Author Only Title Only Author and Title</u>

**Zheng Z, Reichel M, Deveson I, Wong G, Li J, Millar AA (2017) Target RNA Secondary Structure Is a Major Determinant of miR159 Efficacy. Plant Physiol 174: 1764–1778**

Google Scholar: <u>Author Only Title Only Author and Title</u>

**Zhu J-Y, Sae-Seaw J, Wang Z-Y (2013) Brassinosteroid signalling. Development 140: 1615–20**

Google Scholar: <u>Author Only Title Only Author and Title</u>

